# Intensity and dose of neuromuscular electrical stimulation influence sensorimotor cortical excitability

**DOI:** 10.1101/2020.02.21.957928

**Authors:** Ainhoa Insausti-Delgado, Eduardo López-Larraz, Jason Omedes, Ander Ramos-Murguialday

## Abstract

Neuromuscular electrical stimulation (NMES) of the peripheral nervous system has been largely used in the field of neurorehabilitation to decrease muscle atrophy and to restore motor function in paralyzed patients. The rehabilitative effects of NMES rely on the direct or indirect efferent effect on muscle tone and afferent volleys that induce cortical excitation. Although different neuroimaging tools suggested the capability of NMES to regulate the excitability of sensorimotor cortex and corticospinal circuits, to date how intensity and dose of NMES can neuromodulate the brain oscillatory activity measured with electroencephalography (EEG) is yet to be clarified. In the present study, we quantify the effect of NMES parameters on brain oscillatory activity of twelve healthy participants who underwent stimulation of wrist extensors during rest while EEG was recorded. Three different NMES intensities were included: (1) low, inducing slight sensory perception, (2) medium, inducing moderate sensory perception, and (3) high, generating a functional movement. Firstly, we efficiently removed stimulation artifacts from the sensorimotor brain oscillatory activity. Secondly, we analyzed the effect of amplitude and dose on the latter. On the one hand, we observed significant NMES amplitude-dependent brain SMR modulation, demonstrating the direct effect of afferent receptors recruitment. On the other hand, our results revealed a significant NMES amplitude-based dose-effect on SMR modulation over time. While at low and medium intensities the NMES produced a significant cortical inhibitory effect in time, at high intensity a significant cortical facilitatory effect was induced. These results highlight the functionally relevant role of muscle contraction and proprioception in sensorimotor processes, which should be carefully considered for the design and development of NMES based neuromodulation.

## 1. Introduction

Electrical stimulation of the peripheral nervous system is a technique that has been used to study and explore neurophysiology since Luigi Galvani experimented on frogs in 1780 in Bologna. More than two centuries of electrophysiological work resulted in what today are standard clinical measures that allow empiric assessment of nervous system function at different levels (Robinson, 2008). Brain stimulation has been used to map brain-to-muscle or -to-spine connectivity, and peripheral stimulation to map muscle or peripheral sensory receptors (mechanoreceptors, nociceptors, etc.) connectivity to the spine and brain, inducing reflexes and/or brain sensory evoked responses (Rossini et al., 2015). These measurements are used to determine the functioning of different neural networks (e.g., Hoffmann reflex) and can reflect synaptic efficacy at a system level (Pierrot-Deseilligny and Burke, 2005).

Neuromuscular electrical stimulation (NMES) is a peripheral electrophysiological technique that consists of applying electrical currents on the skin to depolarize motor and sensory nerves beneath the stimulating electrodes (Bergquist et al., 2011). NMES has been used as a neuroscientific tool to study sensorimotor neural mechanisms and structures (Carson and Buick, 2019), and also as a clinical application, e.g., in neurorehabilitation to reduce muscle atrophy, and to improve muscle tone and motor function in patients with paralysis after stroke (Knutson et al., 2015; Yang et al., 2019) or spinal cord injury (SCI) (Patil et al., 2015). The working principle of rehabilitative NMES is based on: 1) the direct effect on muscle tone; and 2) the activation of receptors that generate afferent volleys that induce cortical excitation. The expected neuromodulation is based on the pioneering work from Fetz and Baker (1973), who used operant conditioning to generate patterns on precentral activity units and correlated responses in adjacent cells and contralateral muscles (Fetz and Baker, 1973). Although recent ongoing work using oscillation dependent (closed-loop) electrical stimulation has been investigated in animals (Capogrosso et al., 2016; Nishimura et al., 2013) and humans (van Elswijk et al., 2010; Zrenner et al., 2018) to induce plasticity measured by evoked responses, little is known about the effect of the stimulation parameters and dose on the brain oscillatory activity. Most of those works assume a gain modulation of the incoming stimulus depending on the phase of the ongoing oscillation (van Elswijk et al., 2010), thus a neuro facilitatory/inhibitory effect.

Since Hans Berger first worked with the electroencephalography (EEG), this neuroimaging tool has constituted the standard technique to study brain oscillatory activity, with particular focus on sensorimotor processes (including sensory evoked potentials and rhythms) (Birbaumer et al., 1990; Buzsaki, 2006; Shibasaki and Hallett, 2006). The sensorimotor rhythm (SMR) or oscillatory activity, mainly comprised by alpha ([7-13] Hz) and beta ([14-30] Hz) bands, has been thoroughly used to study sensorimotor processes (López-Larraz et al., 2018; Ramos-Murguialday and Birbaumer, 2015; Ray et al., 2019) and can be quantified as the event-related (de)synchronization (ERD/ERS) (Pfurtscheller and Lopes da Silva, 1999). However, to date only a few studies have investigated how NMES modulates the brain sensorimotor oscillatory activity measured by EEG (Corbet et al., 2018; Tu-Chan et al., 2017; Vidaurre et al., 2019, 2016).

Previous work described that brain activation due to NMES, studied by functional magnetic resonance imaging (fMRI) and near infrared spectroscopy (NIRS), is proportional to the applied intensity (Blickenstorfer et al., 2009; Schürholz et al., 2012; Smith et al., 2003). These findings rely on the fact that as the stimulation intensity increases, there is a progressive recruitment of more afferent receptors that modulate brain activity (Golaszewski et al., 2012; Maffiuletti et al., 2008). There is evidence suggesting that below-motor-threshold stimulation activates cutaneous mechanoreceptors that provide feedback to areas 3b and 1 in the somatosensory cortex (S1), while above-motor-threshold stimulation generates muscle contractions that activate muscle spindles and Golgi tendon organs that send afference to areas 3a and 2 in S1 (Carson and Buick, 2019). It is believed that muscle spindles can directly project to the motor cortex (M1) via area 3a, whereas neural transmission due to cutaneous activation from area 3b to M1 is scarce (Carson and Buick, 2019). Thus, the presence or absence of muscle contraction elicited by NMES has a direct impact on somatosensory cortex, and in turn, on motor cortex excitability (Sasaki et al., 2017).

Cortical excitability modulation over corticospinal or corticomuscular connectivity (i.e., over the sensorimotor loop) has been demonstrated using transcranial magnetic stimulation (Fujisawa et al., 2011). Recent experiments investigated the relevant role of peripheral sensory stimulation intensity in influencing corticomotor excitability (Takahashi et al., 2019). While intensities above-motor-threshold induced stronger corticomotor output measured by means of motor evoked potentials (MEP), the results for stimulation below motor threshold led to no agreement, probably due to the wide range of stimulation parameters used in the existing literature (Carson and Buick, 2019; Chipchase et al., 2011). All these results suggest a clear interplay between the cortical activity induced by peripheral stimulation (sensory areas) and the sensorimotor cortical areas with direct corticospinal connections. If some of the sensorimotor cortical connections are preserved, one can assume that corticospinal connectivity could be promoted by peripheral stimulation. This is one of the main arguments for physiotherapy and robot mediated therapies (Alam et al., 2016; Jackson and Zimmermann, 2012) after stroke, especially the ones leveraging closed-loop systems (Mrachacz-kersting et al., 2016; Ramos-Murguialday et al., 2013).

Some interesting dose effects of electromagnetic stimulation in corticospinal connectivity have been also reported during transcranial stimulation (rTMS and tDCS) presenting inhibitory and facilitatory effects due to the number of pulses (i.e., dose) delivered over time (Nitsche and Paulus, 2000; Zrenner et al., 2018). While intensity currents affect the polarization of neuronal membrane facilitating or inhibiting its depolarization, and therefore affecting synaptic efficacy, the effects of prolonged stimulation periods over neural excitability and their consequences on neuroplasticity remain unclear (Knotkova et al., 2019). It has been speculated that the nervous system maintains its excitability within an equilibrium range through homeostatic plasticity adjustments derived from the history of neuronal activity. Therefore, these history- or time-dependent mechanisms can prevent excessive excitatory destabilization reversing its activity towards an opposite state of excitability (Andrews et al., 2013). Similarly, a progressive perceptual adaptation or reduction of sensory responsiveness has been evidenced after prolonged intervals of peripheral vibrotactile and electrocutaneous stimulation (Buma et al., 2007; Graczyk et al., 2018; Leung et al., 2005), indicating a neural compensation after a perturbation of the oscillatory neural system. Furthermore, the capacitance effect present in neural systems (Kiernan et al., 2004; Nodera and Kaji, 2006) could be one of the mechanisms in play during the compensation of the neural oscillatory system. It could be conceivable that after prolonged periods of NMES the sensorimotor cortex could experience similar changes of excitability as result of rebalance of neuronal activity.

In this work we acquired EEG activity from 12 healthy participants during NMES of the wrist extensor muscles at 3 different intensities (2 below and 1 above the motor threshold) in random order to investigate the neuromodulatory effect of peripheral NMES on the ongoing cortical oscillatory activity. We hypothesized that NMES at different intensities would result in the recruitment of distinct afferent receptors and would modulate of the SMR accordingly. Indeed, we speculate that the strongest neuromodulatory changes in the sensorimotor cortex will be achieved with the presence of muscle contractions elicited by high intensity NMES. One might also surmise that the potential of the NMES to regulate the SMR can be dose dependent and exhibit suppression or boosting of excitability over time.

## 2. Materials and methods

### 2.1. Participants

Twelve right-handed healthy participants (four females, age = 27.5±3.0) were recruited to participate in the study. All of them signed an informed consent form. The experimental procedure was approved by the Ethics Committee of the Faculty of Medicine of the University of Tübingen (Germany).

Participants were asked to stay comfortably seated on a chair with their right arm resting on a side table and the hand hanging with the palm facing downwards. Neuromuscular electrical stimulation (NMES) electrodes were placed on the right-hand extensors, as described in Figure 1a. Electroencephalographic (EEG) and electromyographic (EMG) activity was recorded during the experiment. The electrical artifact recorded in the EMG was used to align stimulation onset during the EEG signal processing.

**Figure 1.**
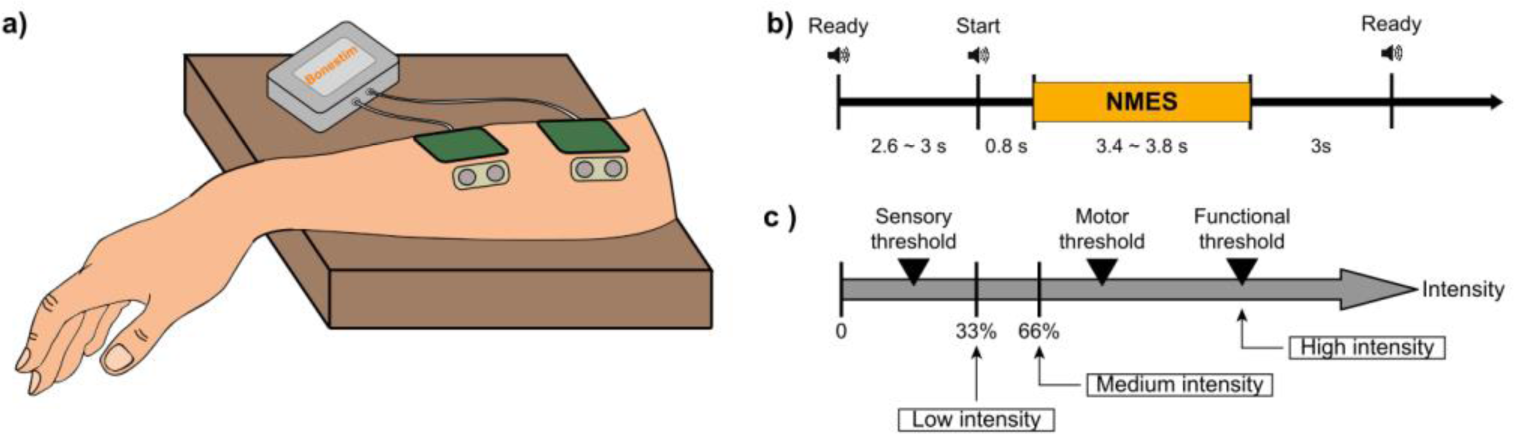
Experimental design and procedure. (a) Representation of the location of stimulation and EMG electrodes placed on right wrist extensors. (b) Timeline of the 3 phases included in each trial: preparation, NMES and inter-trial period. (c) Determination of NMES intensities: low intensity as one third between the sensory threshold and motor threshold, medium intensity as two thirds between the sensory threshold and motor threshold, and high intensity as functional motor threshold.

### 2.2. Experimental design and procedure

The main purpose of the experiment was to investigate NMES neuromodulatory effects (instantaneous and cumulative) on brain oscillatory activity. With this aim, we compared the afferent cortical activity generated by 3 different NMES intensities. Participants were passively stimulated, meaning that they were resting and no volitional motor command was generated during stimulation. Each participant underwent one session consisting of 9 blocks, each comprising 18 trials. One of the three NMES intensities was randomly assigned to each block (determination of the current intensities explained in the section 2.4. Neuromuscular electrical stimulation), resulting in 3 blocks per intensity. A Ready cue was presented 2.6 to 3 seconds before the NMES interval, which lasted between 3.4 and 3.8 seconds. From the offset of the NMES to the next Ready cue, a 3-second inter-trial period was introduced (see Figure 1b). Auditory cues announced the beginning of each interval. The time between blocks was used as breaks, lasting around 150 s (i.e., two and a half minutes). The entire session including setup did not exceed 90 minutes.

### 2.3. Data acquisition

The electroencephalographic (EEG) activity was recorded with a commercial 32-channel Acticap system (BrainProducts GmbH, Germany) and a monopolar amplifier BrainAmp (BrainProducts GmbH, Germany). The recording electrodes were placed at FP1, FP2, F7, F3, Fz, F4, F8, FC3, FC1, FCz, FC2, FC4, C5, C3, C1, Cz, C2, C4, C6, CP5, CP3, CP1, CPz, CP2, CP4, CP6, P7, P3, P4, P8, O1, and O2, following the international 10/20 system. Ground and reference electrodes were placed at AFz and Pz, respectively.

Surface electromyographic (EMG) activity of the right forearm of the participants was recorded by an MR-compatible amplifier BrainAmp (BrainProducts GmbH, Germany) using Ag/AgCl bipolar electrodes (Myotronics-Noromed, Tukwila, Wa, USA) with 2 cm inter-electrode space. Two recording electrodes were placed adjacent to the stimulation pads (see Figure 1a), using the right collarbone as ground. Both EEG and EMG signals were synchronously acquired at a sampling rate of 1000 Hz.

### 2.4. Neuromuscular electrical stimulation (NMES)

A programmable neuromuscular stimulator Bonestim (Tecnalia, Serbia) was used to deliver the stimulation. The cathode (3×3.5 cm, self-adhesive electrode) was placed over the muscles involved in wrist extension (*extensor digitorum* and *extensor carpi ulnaris)*, while the anode (5×5 cm, self-adhesive electrode) was placed 5 cm distal to the cathode. To ensure the correct location of the electrodes, individually determined for each participant, stimulation above motor threshold was applied until a complete wrist extension was induced.

The frequency of the NMES was set to 35 Hz, and the pulse width to 300 µs (Lynch and Popovic, 2008). The individual intensities for each subject were obtained by a scan of currents, starting at 1 mA and increasing in steps of 1 mA. The participants were asked to report the initiation of the following sensations: *(i)* tingling of the forearm (i.e., sensory threshold—STh), *(ii)* twitching of the fingers (i.e., motor threshold—MTh), and *(iii)* complete extension of the wrist (i.e., functional threshold—FTh). According to these thresholds, the three NMES intensities were calculated. Low intensity was defined as one third between the STh and MTh; medium intensity as two thirds between the STh and MTh; and high intensity as the FTh (see Equations 1, 2 and 3 and Figure 1c). None of the participants reported pain or any harmful effect due to the stimulation.

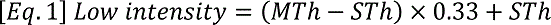

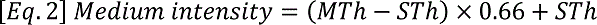

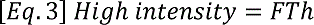

### 2.5. Data preprocessing and analysis

#### 2.5.1. Artifact removal procedures

One important limitation for the quantification of EEG activity during continuous stimulation is the contamination of the signals due to the electrical currents delivered to the body. The EEG is easily polluted by these currents, and artifact removal methodologies are essential to properly estimate cortical activation. With this aim, different techniques for contamination removal in invasive and non-invasive brain activity recordings have been proposed, such as interpolation, blanking or linear regression reference (LRR) (Iturrate et al., 2018; Walter et al., 2012; Young et al., 2018). Blanking of the data is the most restrictive method as contaminated data are rejected and signals that could be of interest are neglected for further analysis. However, if the removal is implemented using hardware, the artifact has less influence on the recovery period of the amplifier preventing it from being saturated and allows the use of other methods to compensate for the missing data (Kent and Grill, 2012). Another approach is to linearly interpolate the corrupted data, connecting the last point before the artifact and the first point after the artifact. However, interpolation induces a bias in the estimation of power spectrum of the signals (Walter et al., 2012). LRR re-references the signals through weights that are assigned to each channel. The weights are calculated in a training block according to the noise of each channel generated by the electrical stimulation. This method effectively reduces artifacts, but it fixes the weights and assumes no changes in channel noise during the intervention. So far, the feasibility of this method has only been proven in invasive recordings (Young et al., 2018), in which impedances are less likely to change within sessions and are more similar among channels (Ball et al., 2009). Normally, impedances deteriorate and noise-influence increases throughout an EEG session complicating the implementation of LRR in non-invasive recordings of brain activity.

Therefore, we implemented an alternative two-step artifact removal method and demonstrated its feasibility. The raw EEG signals were pre-processed using custom-developed scripts in Matlab (MathWorks, Natick, MA, USA).

##### 2.5.1.1. Channel removal based on power-line noise

During an EEG session, particularly during setup and during periods between experimental blocks, special care is required to maintain EEG signal clean (i.e., raw data inspection and impedance check). However, our empirical experience shows that, sometimes, certain EEG electrodes present higher contamination due to the stimulation than others (see Figure 2a). These electrodes present broadband artifacts that impede further analyses even after applying the median filter preprocessing described below. We hypothesized that this effect might be due to degraded impedances, which occasionally deteriorate even when during the setup were set below 5 kOhm. Despite we did not store the impedances of each electrode to check this and discard the electrodes with high impedance, we ideated an automatized method to detect and discard them offline. This way, we could automatically eliminate contaminated channels without the human bias that would constitute a manual rejection.

**Figure 2.**
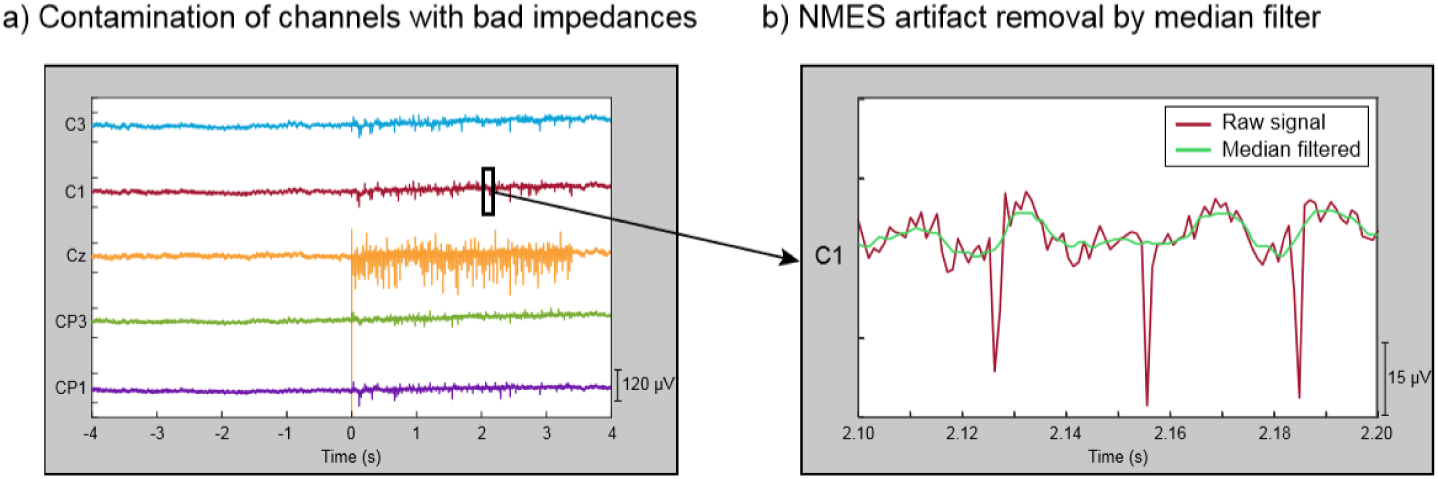
Characterization of contamination induced by NMES. (a) EEG of a representative trial showing non-contaminated channels (C3, C1, CP3, CP1) and one contaminated channel (Cz) during high intensity stimulation. Channels with bad impedances are more prone to be contaminated by electrical stimulation. (b) Zoom in 100 ms segment of a representative EEG trial in a non-contaminated channel that presents NMES artifact (red line). The effect of stimulation artifact is minimized by median filter (green line).

Having high impedance between the recording electrode and the skin resembles an open circuit, where the electrode behaves like an antenna and captures outside electric frequencies (like the power-line noise). Our method exploits this effect and identifies the EEG electrodes with unusually high power-line noise (50 Hz in Europe). Since all the electrodes should be approximately equally exposed to electromagnetic signals at 50 Hz, we assume that very high power at this frequency is an indirect indicator of high skin-electrode impedance. The procedure was applied block-wise, meaning that it was used to detect and remove contaminated channels within each individual EEG block. The EEG activity was high-pass filtered at 0.1 Hz with a 4^th^ order Butterworth. The power spectral distribution at 48-52 Hz was estimated using Welch’s method, averaging the periodogram of 1-second Hamming windows with 50% overlapping. The power mean and standard deviation (SD) of all the EEG channels were calculated from all trials in each block. Channels whose power was higher than 4 SD above the mean were discarded from that specific block. The remaining channels were used to re-compute the mean and SD. The procedure was iteratively repeated until no channels exceeded the rejection threshold (see Figure 3 dashed box, Supplementary Figure 1 and Supplementary Figure 2).

**Figure 3.**
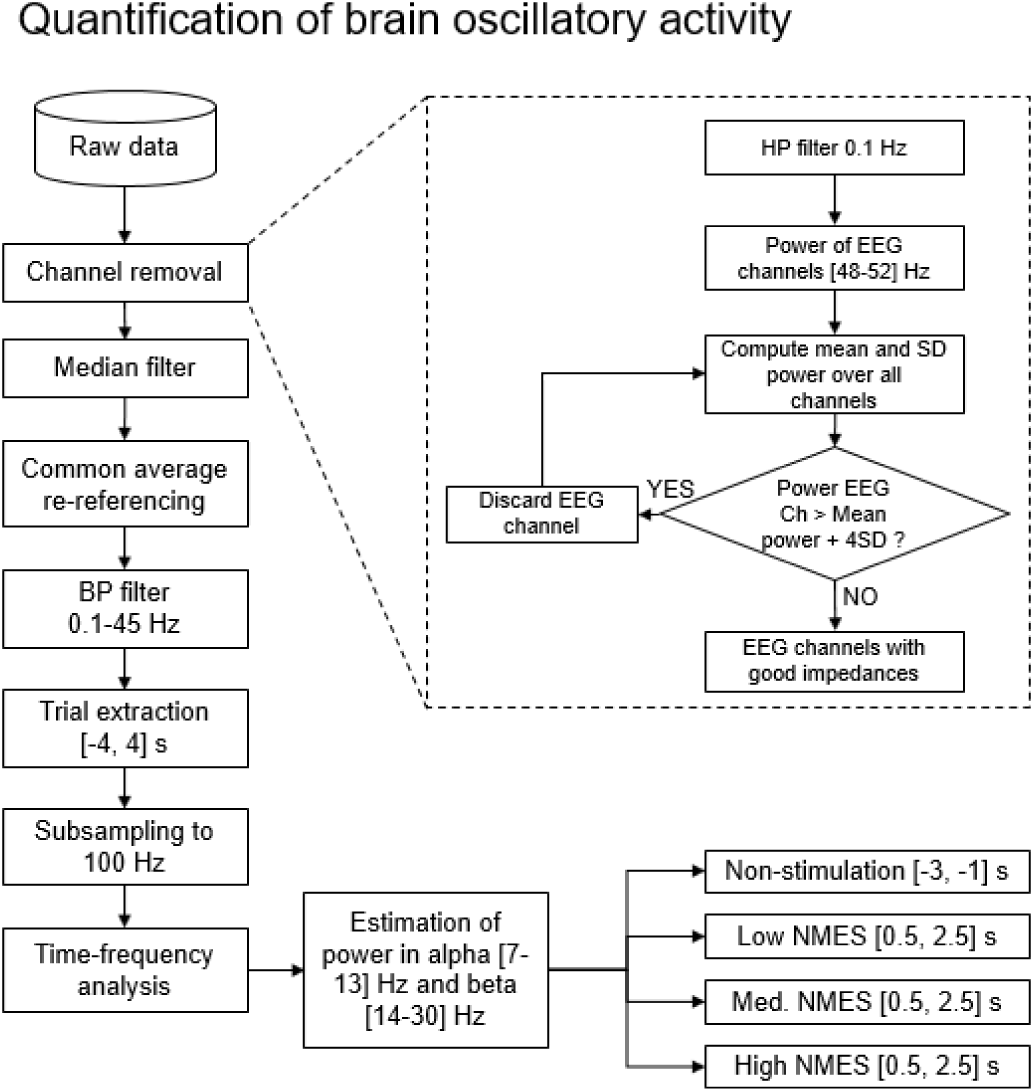
Flowchart with the steps for the quantification of brain oscillatory activity. The whole set of data is firstly preprocessed by a two-step procedure based on channel removal and median filtering. In this level, channels with bad impedances and artifacts due to electrical stimulation are removed. Then, the remaining clean data is filtered and divided into trials. Finally, power is estimated in alpha [7-13] Hz and beta [14-30] Hz bands for non-stimulation ([-3, −1] s) and NMES ([0.5, 2.5] s) intervals, being the baseline [-2.5, −1.5] s.

##### 2.5.1.2. Median filtering for removal of electrical stimulation contamination

It is well known that applying electro-magnetic currents to stimulate the neural system can introduce undesired noise to the recordings. The NMES configuration used in this study introduces large peaks of short latency (∼5 ms) to the recorded EEG and EMG signals. Therefore, median filtering was used to minimize the NMES induced artifacts (Insausti-Delgado et al., 2017). This filter is suited to eliminate high-amplitude peaks from a time series (Gallagher and Wise, 1981), and can remove the short-latency high-amplitude artifacts caused by the NMES. A sliding window of 10 ms was applied to the EEG signal in steps of one sample, providing as output the median value of each window. We selected a 10 ms window as it fully covers the electrical artifact. This filter produces a frequency-dependent attenuation that follows an exponential function from 0 to 100 Hz (i.e., the frequency with a period that completely fits within the 10 ms window), leading to low attenuation at low frequencies and a complete attenuation at 100 Hz (Supplementary Figure 3). With this window size, the attenuation of the signal at 10 Hz, 20 Hz and 30 Hz is 1.28%, 4.89% and 10.90%, respectively. The relatively low attenuation at low frequencies makes this method suitable for analyzing alpha and beta sensorimotor oscillations. Figure 2b displays a zoomed segment of 100 ms of activity, showing the effect of the median filter on the stimulation artifacts.

### 2.6. Quantification of brain oscillatory activity

A common average reference (CAR) was applied to the EEG signals. The re-referenced signals were band-pass filtered at 0.1-45 Hz using a 1^st^ order Butterworth filter. Each block was trimmed down to (18 x) 8-second trials, from −4 seconds to +4 seconds, being 0 the beginning of the stimulation. The trials were down sampled at 100 Hz, and those belonging to the same level of NMES intensity were pooled together.

The quantification of cortical activity was performed by evaluating the spectrum differences of the sensorimotor rhythms, by means of the alpha and beta event-related (de)synchronization (ERD/ERS), i.e. decrease or increase in power generated by an event compared to a baseline (Pfurtscheller and Lopes da Silva, 1999). Large ERD values (i.e., more negative power values) represent stronger cortical activation compared to baseline time interval, as it represents disinhibition/excitation of neural population activity (Ritter et al., 2009). For the quantification of brain activity, we used the Fieldtrip toolbox (http://fieldtriptoolbox.org/) for Matlab. Time-frequency maps were calculated using Morlet wavelets in the frequency range from 1 to 45 Hz, with a resolution of 0.5 Hz. The power change was computed as the percentage of increase or decrease in power (i.e., ERS or ERD) with respect to the baseline ([-2.5, −1.5] s), as described in Equation 4.

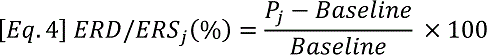

From the time-frequency maps, we calculated the mean change in power of channels C1, C3, CP1 and CP3 (i.e., area over the sensorimotor cortex representing right forearm, contralateral hemisphere to the stimulated limb), since we considered that averaged values over these electrodes could better quantify the overall changes in the sensorimotor areas. We calculated the sensorimotor averaged changes in power in alpha [7-13] Hz and beta [14-30] Hz bands for non-stimulation ([−3, −1] s) and NMES ([0.5, 2.5] s) intervals, using as baseline the [-2.5, −1.5] s interval (see Figure 3). The NMES period was defined as starting at 0.5 s to avoid potential bias and influence of the stimulation onset like event-related brain potentials (e.g., error potentials, P300, etc.) at t=0. For EEG topographical inspection and analysis, all channels were analyzed individually.

### 2.7. Statistical analysis

The statistical tests were performed in IBM SPSS 25.0 Statistics software (SPSS Inc., Chicago, IL, USA) and Matlab. We used the Shapiro-Wilk test to determine the normality of the data. Accordingly, a multivariate analysis of variance (MANOVA) for repeated measures was performed to find differences in the dependent variables, alpha and beta ERD/ERS, with NMES intensity (4 levels: no stimulation, low, medium and high intensity stimulation) as within-subject factor. In order to determine the origin of the significant effect, post-hoc tests with Bonferroni correction were performed. In order to analyze whether NMES can induce a dose-effect, we studied the ERD/ERS changes over time. For that, we computed the alpha and beta ERD/ERS for each single trial (i.e., in Eq. 4, *Pj* was the alpha/beta power of each trial during the NMES period, and the baseline was calculated from the grand average of all the trials of each intensity). A linear regression was estimated for the ERD/ERS values over trials for the two frequency bands (i.e., alpha and beta) and the three NMES intensities (i.e., low, medium and high). Correlation between ERD/ERS and sequence of trials were calculated using Pearson’s correlation coefficient to study stimulation effects over time.

## 3. Results

### 3.1. Effect of artifact removal

The pre-processing of the data eliminated satisfactorily the electrical noise contamination coming from the peripheral electrical stimulation. It reduced the effect of the artifacts to an extent that allowed us to perform EEG spectral analysis of the brain oscillatory activity.

Channels with good impedances are also influenced by the electrical stimulation artifact that is introduced into the signal as large peaks. Figure 2b illustrates in a 100 ms segment of a representative trial how the median filter deals with these undesired artifacts. To prove the efficacy of the method, we focused on the worst-case scenario, as the contamination is larger for higher stimulation intensities. This effect can be observed in Figure 4, which depicts the EEG time-frequency activity at the different NMES intensities including artifacts, and after the median and spatial filters are applied. The NMES generates an increase of power, or ERS, around 35 Hz (i.e., the stimulation frequency), which increases as the NMES intensity is incremented (Figure 4, left column). This power increase in high-beta/low-gamma band due to stimulation artifact was eliminated for all intensities after median filtering. Applying the median filter did not change the power in alpha and beta frequencies before the stimulation onset (t = 0 s), but eliminated the ERS during the stimulation period, minimizing the artifacts and revealing the alpha and beta modulation. The common average re-reference (CAR) after median filtering enhanced the power decrease of the bands of interest for every NMES intensity, as it does with non-contaminated EEG (Wolpaw et al., 2002). Regardless of the intensity delivered, the stimulation generated the classical event-related potentials (e.g., error potentials, P300, etc. (Duncan et al., 2009)) between −0.8 and 0.5 s, due to the cue presentation (−0.8 s) and especially due to the sensory perception of the NMES initiation. This can be seen as power increase at low frequencies (1-5 Hz).

**Figure 4.**
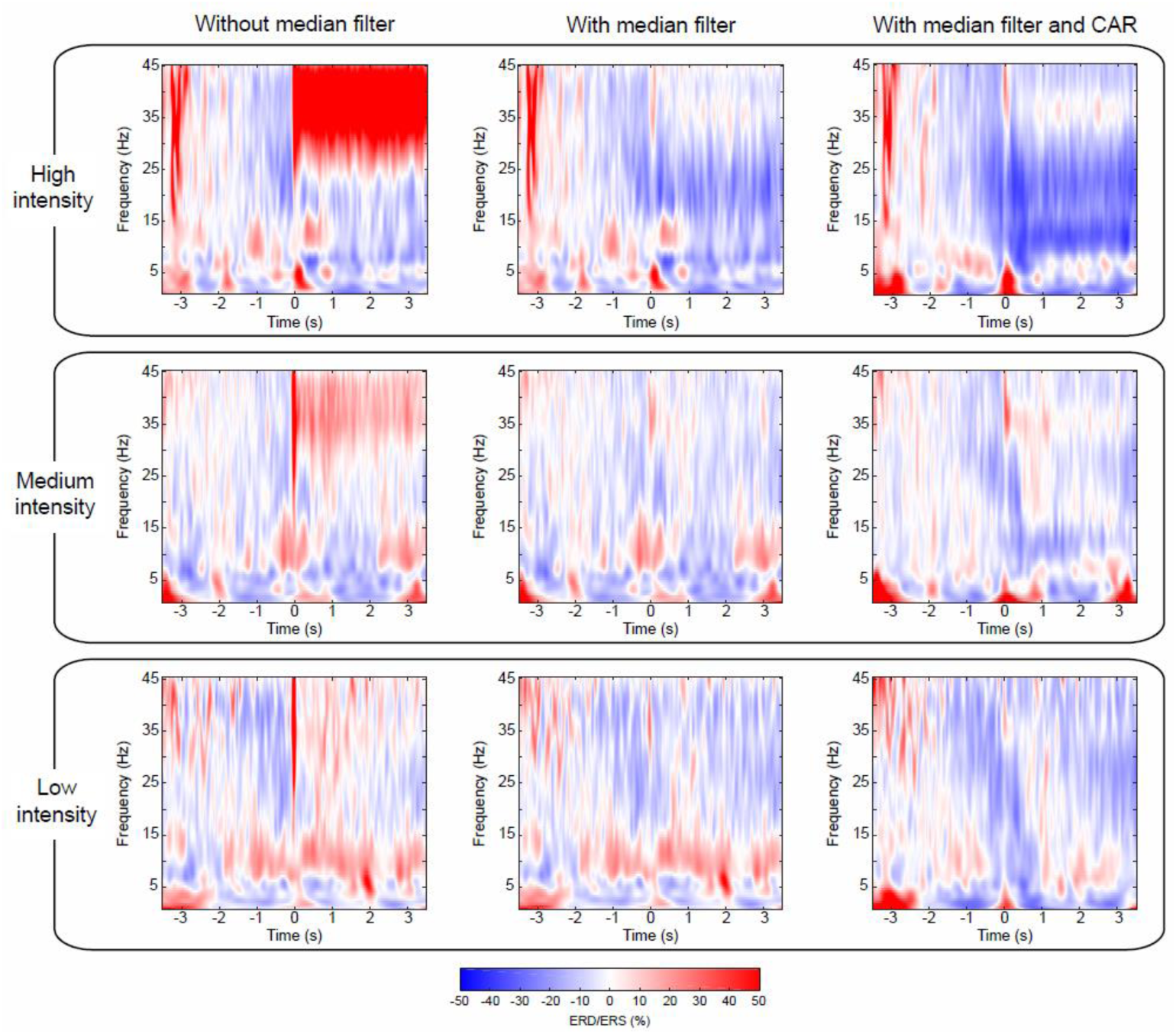
Comparison of cortical activation after median and spatial filtering. Time-frequency maps averaged over all participants, representing ERD/ERS, of the average of channels (C3, C1, CP3, CP1) located over the contralateral motor cortex to the stimulated limb. Averaged time-frequency maps without median filter (left column), with median filter (center column), and with median and CAR filter (right column) after removal of contaminated channels. Panels show the different NMES intensities: high (top panel), medium (middle panel) and low intensity (bottom panel). The percentage of ERD/ERS is computed according to the baseline [-2.5, −1.5] s. Time 0 s is aligned with the onset of the stimulation.

### 3.2. Influence of stimulation intensity on cortical activation

To analyze the influence of stimulation intensity on cortical activation, we compared the changes in brain oscillatory activity in four conditions: non-stimulation, low, medium and high intensity stimulation (see topoplots of alpha and beta rhythms in Figure 5). Topographic maps of non-stimulation condition were calculated using the interval [-3, −1] s prior to the stimulation, while the other conditions were extracted from the interval [0.5, 2.5] s after stimulation onset. An increment of the stimulation intensity resulted in an increasing ERD (i.e. larger decrease in power) in both frequency bands over the sensorimotor cortex as expected (Blickenstorfer et al., 2009; Schürholz et al., 2012; Smith et al., 2003), while occipital areas showed idling activity. At high intensity stimulation sensorimotor cortex of both hemispheres presented a decrease of power, being more pronounced in the contralateral hemisphere, as demonstrated in previous work studying brain oscillatory signatures of motor tasks (Ramos-Murguialday and Birbaumer, 2015).

**Figure 5.**
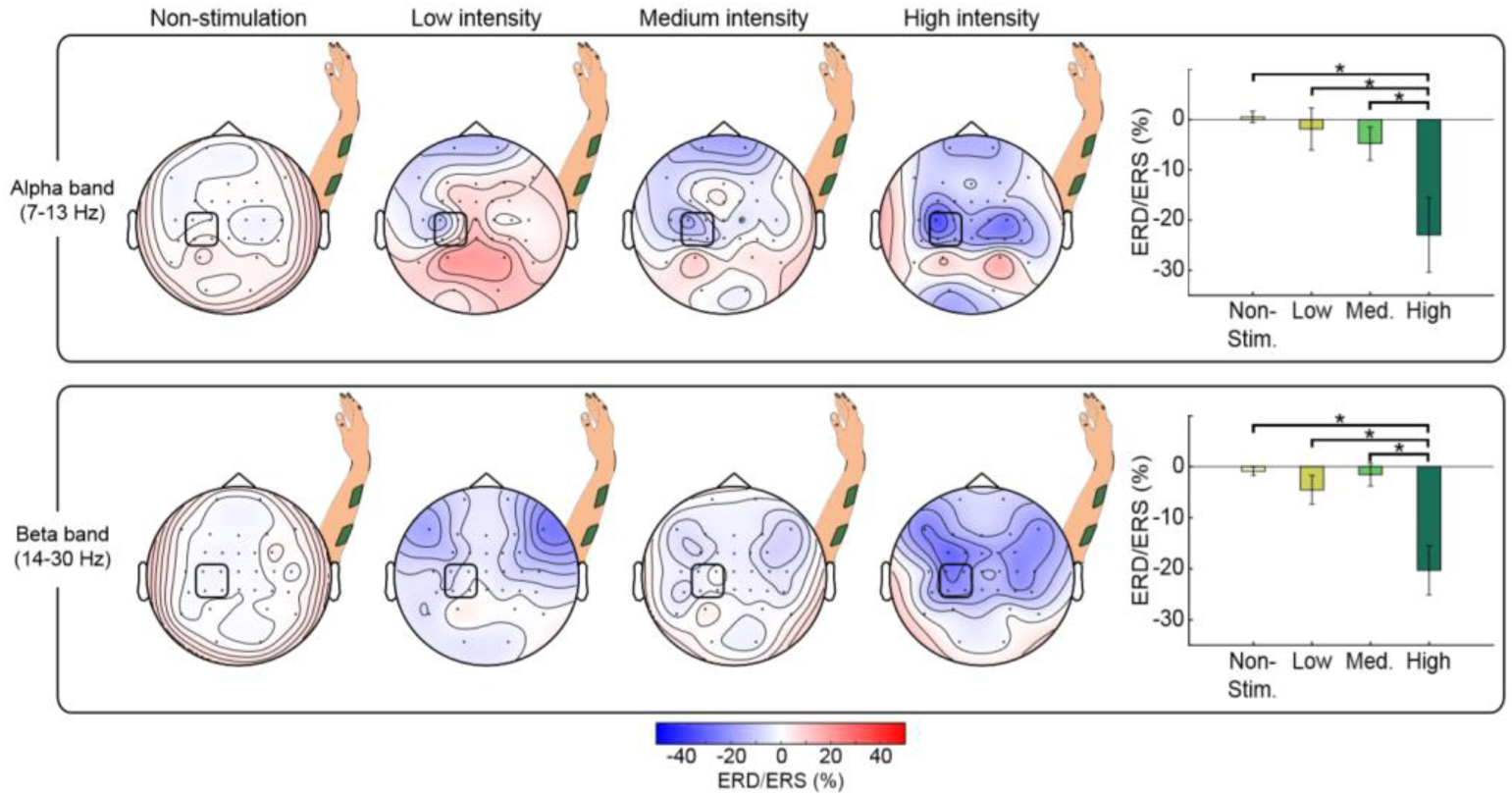
Comparison of cortical activation for different stimulation intensities for alpha and beta band EEG activity. Topographic maps, averaged over all participants, showing ERD/ERS of non-stimulation periods [-3, −1] s and NMES periods [0.5, 2.5] s belonging to each intensity (i.e., low, medium and high) for alpha (upper row) and beta (lower row) frequency bands. Bar graphs show the mean percentage of ERD/ERS averaged from channels (C3, C1, CP3, CP1) for each intensity and frequency band. The statistically significant differences between pairs are expressed with horizontal lines and stars. The percentage of ERD/ERS is calculated with respect to the baseline [-2.5, −1.5] s. The signals are processed using the two-step procedure (i.e., removal of contaminated channels and median filter) and a CAR.

Our MANOVA analysis, reflected a significant effect of intensity on alpha and beta ERD (*F*(6, 86) = 4.356, *p* = 0.001). Rightmost panels in Figure 5 display the results of the post-hoc comparisons. For both alpha and beta, there was a significantly higher ERD (i.e., more negative values) induced by high intensity NMES compared to rest (*p* = 0.004 for alpha, *p* < 0.001 for beta), to low intensity NMES (*p* = 0.013 for alpha, *p* = 0.004 for beta) and to medium intensity (*p* = 0.045 for alpha, *p* < 0.001 for beta).

### 3.3. Stimulation dose-effect

To study the influence of stimulation dose on cortical activity, we computed the ERD/ERS of each single trial and performed a regression in time within each block (trials were performed sequentially) and within session appending same stimulation intensity blocks in order of appearance (blocks of different intensities were presented semi-randomly, i.e., most likely after low intensity block a high intensity or medium block might have been presented). Figure 6 shows the average ERD/ERS for all participants in both frequency bands during the 54 trials (3 blocks x 18 trials, see vertical yellow lines) for each intensity and a linear regression to fit them.

**Figure 6.**
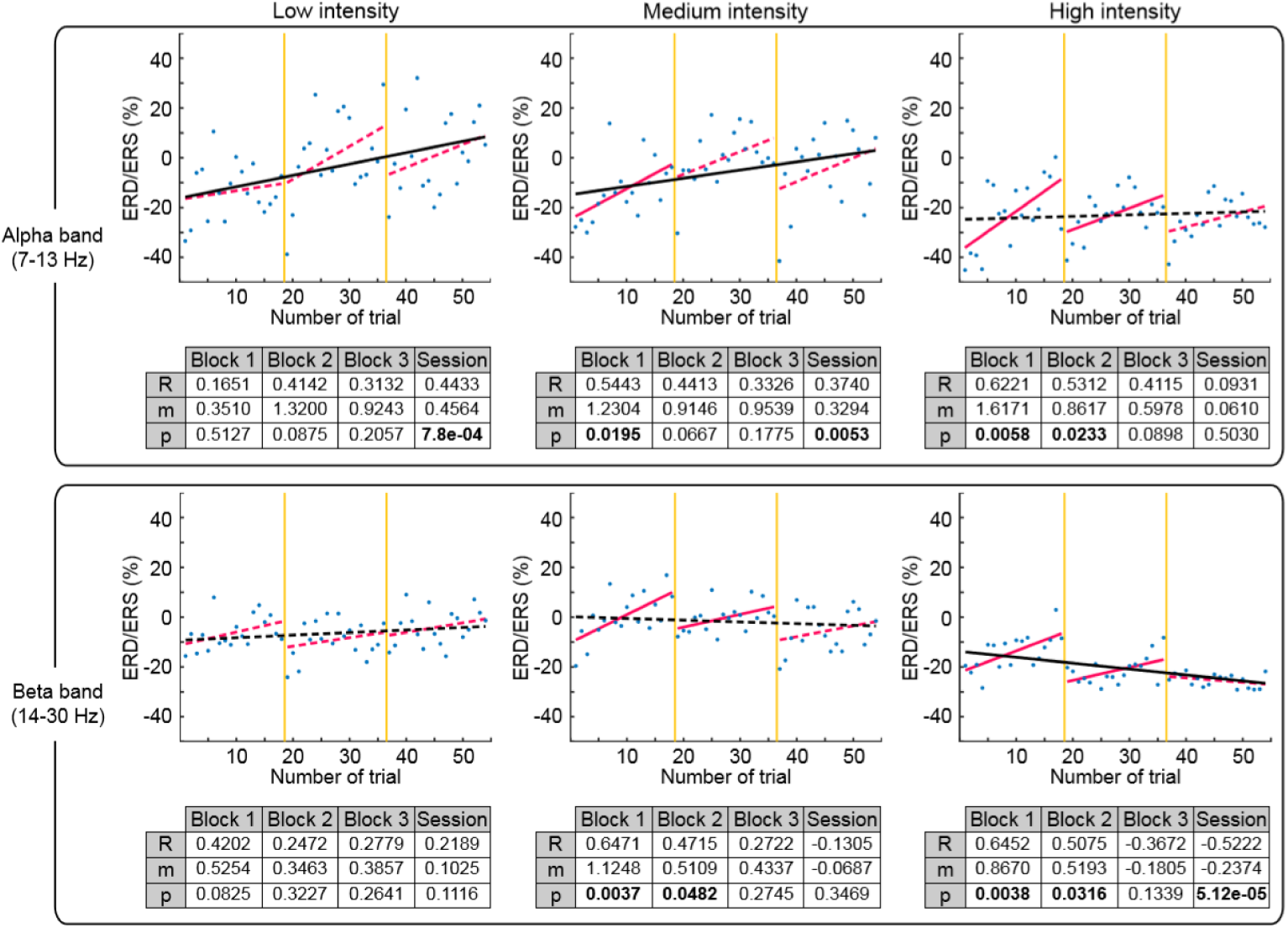
Comparison of cortical activation over trials for alpha and beta band. The cortical activity during NMES period ([0.5, 2.5] s) quantified as ERD/ERS over the 54 trials divided into blocks by vertical yellow lines of each stimulation intensity, averaged for all the participants. The percentage of ERD/ERS is calculated according to the baseline [-2.5, −1.5] s. Different intensities are compared in columns: low (left), medium (middle) and high (right). Alpha (upper row) and beta (bottom row) frequency bands are described. Significant correlation between ERD/ERS and sequence of trials over session are represented with black solid linear regressions. Within block significant correlations are displayed by magenta solid linear regressions. Tables show the correlation (R), slope (m) and p-value (p) for every block and session.

The first thing we observed is a clear modulation based on stimulation, reducing its variability with stimulation amplitude. Low and medium intensities caused a significant reduction of alpha ERD over time (*p* = 7.8e-4 for low; *p* = 0.0053 for medium). In contrast, high intensity caused a significant enhancement of beta ERD over time (*p* = 5.12e-5). Furthermore, as can be seen in Figure 6, we observed that in every block (magenta lines separated by yellow vertical lines) there is a reduction of the ERD (increase of power plotted as linear regression) progressively induced per trial from the first to the last trial. The first trial of the new block presented a larger ERD (decrease of power) in comparison to the ERD of the last trial of the previous block (irrespective of the stimulation intensity and order of the block within the session).

## 4. Discussion

This study demonstrated the significant effects of artifacts, intensity and dose of neuromuscular electrical stimulation (NMES) of the upper limb on the ongoing brain oscillatory activity recorded using EEG.

First of all, we dealt with the issue of artifact removal to allow accurately estimating the cortical oscillatory activity. Recordings of brain activity are easily polluted, especially when electrical stimulation interacting with the nervous system is concurrently used. This contamination can negatively affect the signal to noise ratio, covering the brain activity. Our findings evidenced that the median filter enhanced the detection of sensorimotor oscillatory activity after removing stimulation artifacts.

After the EEG data was cleaned, especially of NMES-induced artifacts, we analyzed the modulation of alpha and beta oscillations produced by the stimulation. Power suppression, or desynchronization, of these frequencies has been associated with cortical excitation, whereas synchronization reflects a state of inhibition (Klimesch et al., 2007). During high intensity NMES, the induced desynchronization in alpha and beta was significantly larger than during stimulation at low or medium intensities or no stimulation. While below motor threshold stimulation intensities only activate cutaneous mechanoreceptors (e.g., Pacinian corpuscles and Merkel disks), stimulation above motor threshold also recruits proprioceptive receptors (e.g., muscle spindles, Golgi tendon organs and joint afferents) (Golaszewski et al., 2012; Maffiuletti et al., 2008). It has been proposed that muscle spindles can directly influence the motor cortex (M1) through the area 3a, while the projections from area 3b activated by cutaneous feedback to M1 are less likely to happen (Carson and Buick, 2019). We can therefore assume that high intensity NMES leads to higher cortical excitation, probably by recruiting larger number of receptors derived from muscle contractions in addition to cutaneous afference that is also engaged in sensory-threshold stimulation, and that the recruitment of muscle spindles results in activation of M1 via area 3a of the S1 (Carson and Buick, 2019; Schabrun et al., 2012). These results of intensity dependent brain activation are in line with corticomuscular responses (Sasaki et al., 2017), metabolic responses recorded by functional magnetic resonance imaging (fMRI) and near infrared spectroscopy (NIRS), which demonstrated a direct quantitative association with stimulation intensity (Blickenstorfer et al., 2009; Schürholz et al., 2012; Smith et al., 2003). Noteworthy, our results showed that the cortical activity during low and medium intensities activating only cutaneous mechanoreceptors was not significantly different to no stimulation. This suggests that although there might be an influence of the peripheral stimulation on the ongoing brain oscillatory activity, when there is no muscle contraction and proprioception (due to the movement), the afferent activity reaching the brain does not induce significant cortical modulation measurable in the EEG at the analyzed frequencies.

It is well known that during voluntary movement, a stronger cortical activation is seen in alpha than in beta (López-Larraz et al., 2014; Ramos-Murguialday and Birbaumer, 2015). Such modulation in alpha and beta cortical activity has been related to control top-down and bottom-up neural processes, suggesting its role in the integration of motor tasks preparation and execution with movement-related sensory feedback. However, this balance between alpha and beta rhythms is altered in absence of top-down regulation. Passive mobilizations (e.g., bottom-up transmission) exhibit stronger beta band activity compared with active movements, indicating the relationship of this frequency band with proprioception without volitional muscle contraction (Alegre et al., 2002; Ramos-Murguialday and Birbaumer, 2015), and thus the inhibitory effect of top-down neural control in this frequency. In this study, the peripheral electrical stimulation at the functional intensity level not only generated passive movement of the limb, but also non-volitional contraction of the forearm muscles. This stimulation intensity induced more significant brain activation in beta than in alpha frequency band, suggesting that changes in beta oscillatory activity comprises two components: proprioception and afference of muscle contraction (without volition) through Golgi tendons and muscle spindles. Therefore, our results give a hint of the relevance of beta rhythm on bottom-up neuromodulation (i.e., afferent cortical excitation). In agreement with previous studies, we can speculate that there might be differences in the mechanisms of afferent modulation between passive movements and functional electrical stimulation. Whereas NMES over the motor threshold recruits afferent axons from muscle spindles, Golgi tendon organs and cutaneous receptors (Bergquist et al., 2011; Golaszewski, 2017), Golgi tendon organs are less sensitive to passive movements and discharge less (Paillard and Brouchon, 1968; Purves et al., 2004), and the firing rate of the muscle spindles is muscle lengthening dependent (Chye et al., 2010). However, it cannot be concluded whether our functional NMES induces stronger beta activity than passive movement since we did not include the latter condition in our experimental protocol.

We tracked changes of the SMR and evidenced that NMES induced a dose-effect on brain oscillatory activity over time. Regardless the stimulation intensity applied, both alpha and beta bands presented a short-term reduction of ERD (reduction of brain excitatory effect of NMES) between consecutive trials within a block, showing that the ERD response of a specific trial depended on the previous stimulation. This reduction of ERD vanished at the beginning of every new block, demonstrating the ability of the SMR to reset its excitability after the ≈2-minute inter-block period. However, the overall activity throughout the session depicts that long-term effects survive temporary resets and exhibits a dose-effect over time, suggesting a conditioning effect. We observed different long-term modulatory responses between SMR in alpha and beta bands conditioned by the stimulation intensity. For sensory-threshold intensities (i.e., low and medium), the power in alpha band was significantly reduced throughout the session, suggesting a habituation effect (Leung et al., 2005). NMES at sensory-threshold recruits cutaneous receptors (without eliciting any muscle contraction or movement) that provide sensory afference to the brain (Maffiuletti et al., 2008). It has been evidenced the relevance of alpha band in information processing of attention and awareness (Händel et al., 2011; Klimesch, 2012), and we speculate that the repeated activation of functionally irrelevant sensory afference (i.e., cutaneous afference in absence of movement) results in inhibition of the alpha SMR. A progressive habituation or desensitization (i.e., reduction of perceived sensation) of sensory perception is also presented after prolonged vibrotactile and electrocutaneous stimulation (Graczyk et al., 2018), which is slower at high stimulation intensities (Buma et al., 2007). This desensitization might be caused by a hyperpolarization of axon membranes (i.e., increasing membrane inhibition) controlled by the activity of Na-K pump that prevents the membrane from excessive excitation due to the repetitive electrical stimulation (Kiernan et al., 2004; Nodera and Kaji, 2006). One can hypothesize that the desensitization of sensory perception and habituation of alpha band might be connected somehow.

The modulation of beta power due to stimulation at functional-threshold intensity incremented with time, indicating an excitatory effect on the sensorimotor neural network. Beta oscillations have been related to the neural transmission from the primary motor cortex to the muscles and back to the motor cortex, via afferent pathways and somatosensory cortex (Aumann and Prut, 2015; Khademi et al., 2018). This closed-loop neural network provides the sensorimotor cortex with information of movements, comprising the muscles and joints. NMES at functional intensity recruits proprioceptive receptors (e.g., muscle spindles, Golgi tendon organs and joint afferents) in addition to cutaneous mechanoreceptors (Golaszewski et al., 2012; Maffiuletti et al., 2008). The activation of this larger number of receptors keeps the aforementioned loop working and results in higher excitability of the network over time represented. Humans are constantly preforming motor tasks and movement drives our behavior and has driven our nervous system development. We can hypothesize that only functionally relevant afferent information excites the sensorimotor cortex, while afferent information not related to movement (i.e., mechanoreception due to low and medium NMES) is not considered as “relevant” and is neglected suppressing cortical excitability (Schabrun et al., 2012). All this highlights the different effect when muscle contraction and proprioception enrich afferent activity (probably due to their ecologically relevant role in sensorimotor function), indicating that habituation or attention shift during a movement is less likely to occur due to its functional role in sensorimotor function.

To the best of our knowledge, this is the first time a sensorimotor cortical facilitation and inhibition effect due to NMES has been measured using EEG and characterized as presenting significant intensity- and dose-effects, which occurred in a short period of time. Understanding how NMES parameters, such as intensity and dose, can modulate the excitability of cortical oscillations will allow a better understanding of peripheral electrical stimulation sensorimotor integration. Further work should disclose whether more functional afferent activity (including proprioception and muscle contraction) is needed to increase functional plasticity and modulate sensorimotor function (e.g., corticomuscular synaptic efficacy, cortico-cortico functional connectivity, etc.). The effect of ongoing activity and other stimulation parameters (i.e., pulse width, pulse form, frequency and energy) need to be carefully studied, probably based on computational neuroscience and bioelectromagnetic modeling, to understand their effect in excitatory and inhibitory mechanisms. Nevertheless, the presented results shed some light onto the neuromodulatory mechanisms that can be investigated and exploited using NMES.

## Acknowledgements

This study was funded by the Bundesministerium für Bildung und Forschung BMBF MOTORBIC (FKZ 13GW0053) and AMORSA (FKZ 16SV7754), the Deutsche Forschungsgemeinschaft (DFG), and the Fortüne-Program of the University of Tübingen (2422-0-1 and 2556-0-0 to ELL, and 2452-0-0 to ARM). The work of AID was funded by the Basque Government’s scholarship for predoctoral students.

## Supplementary materials

**Supplementary Figure 1.**
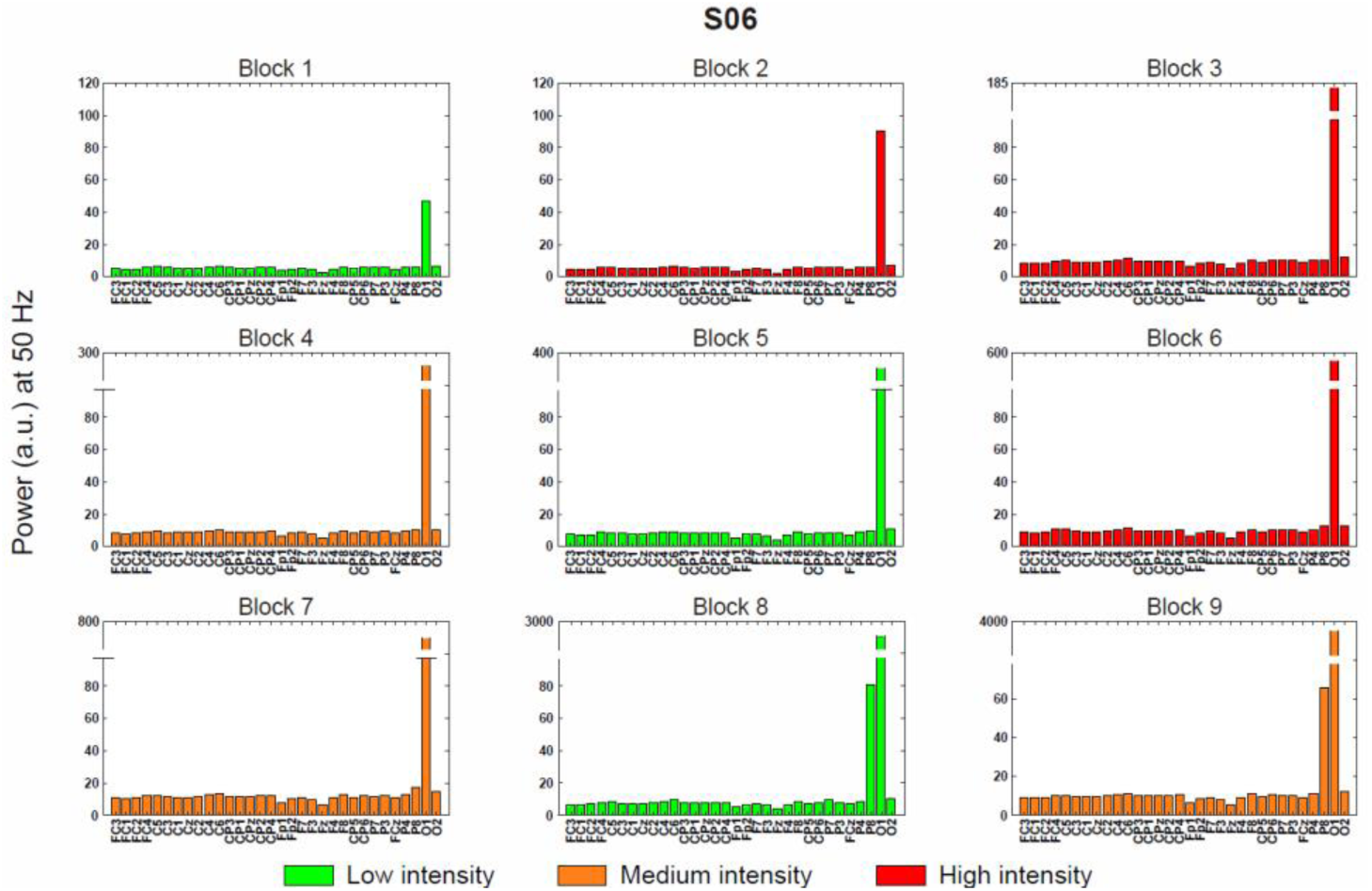
Estimation of power-line noise for channel removal. The barplots represent the power-line noise in each EEG channel for every block of a representative subject. The green, orange and red color of the bars is associated with low, medium and high NMES intensity, respectively. High power-line noise was as an indicator of bad impedance and bad electrode-skin conductivity. We proposed a method based on the removal of the channels that exceeded the mean EEG power + 4 standard deviations. In this particular subject, O1 was rejected from all the blocks, and P8 from block 8 and block 9.

**Supplementary Figure 2.**
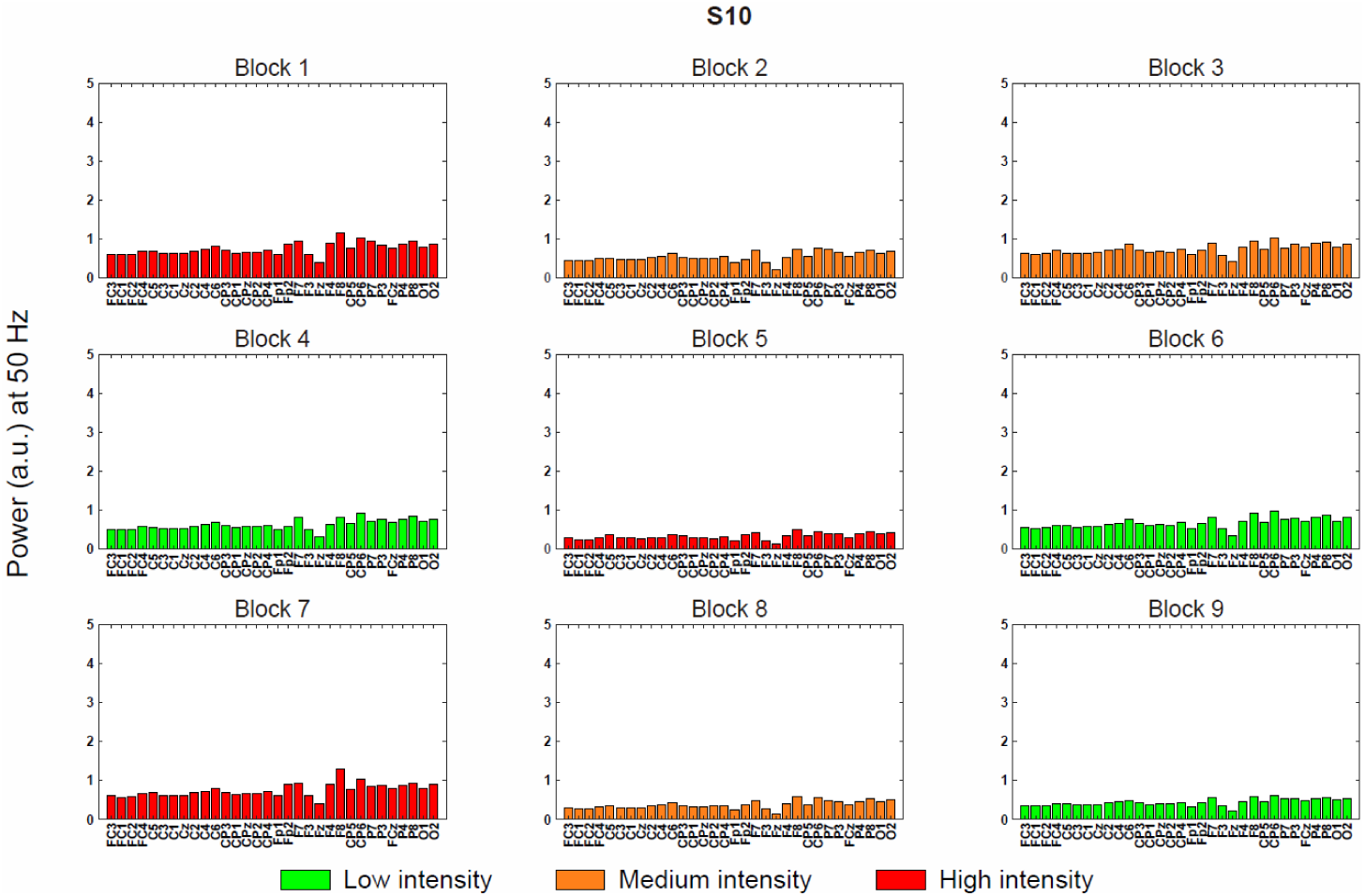
Estimation of power-line noise for channel removal. The barplots represent the power-line noise in each EEG channel for every block of a representative subject. The green, orange and red color of the bars is associated with low, medium and high NMES intensity, respectively. High power-line noise was as an indicator of bad impedance and bad electrode-skin conductivity. We proposed a method based on the removal of the channels that exceeded the mean EEG power + 4 standard deviations. In this particular subject, all the channels in every block satisfied the inclusion criteria.

**Supplementary Figure 3.**
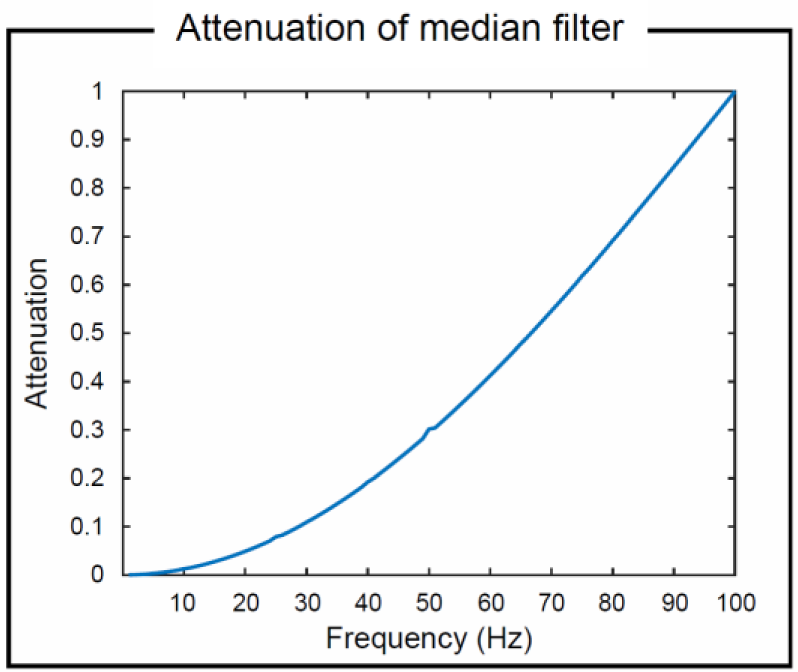
Attenuation of median filter. The median filter induces an exponential attenuation of the signal, which is frequency dependent. This filter attenuates from 0 (no attenuation) to the inverse of the selected window length of the filter; in our case the window of 10 ms produces a total attenuation at 100 Hz. As an example, the attenuation at 10 Hz, 20 Hz and 30 Hz is 1.28%, 4.89% and 10.90%, respectively.

